# Deciphering chromatin architecture and dynamics in *Plasmodium falciparum* using the nucDetective pipeline

**DOI:** 10.1101/2025.08.13.670116

**Authors:** Simon Holzinger, Leo Schmutterer, Victoria Marie Rothe, Maria Theresia Watzlowik, Uwe Schwartz, Gernot Längst

## Abstract

High-resolution analysis of cellular chromatin structure is crucial for uncovering developmental and cell-type-specific regulatory networks. We developed the nucDetective pipeline to provide a comprehensive evaluation of chromatin organisation. This involves assessing nucleosome positioning, occupancy, fuzziness, and array regularity. The pipeline was benchmarked by analysing the chromatin structure of the malaria-causing parasite *Plasmodium falciparum* (*Pf*) during its erythrocytic development cycle. *Pf* is characterised by a unique chromatin landscape, exhibiting unstable nucleosomes and a genomic AT-content exceeding 80%, which presents challenges for standard MNase-seq analysis of chromatin. The nucDetective pipeline provides specific, high-resolution nucleosome profiles for the different asexual stages of *Pf*, monitoring the dynamics of individual nucleosomes. In contrast to the current view, suggesting an irregular chromatin structure and lack of nucleosomal DNA, we demonstrate that the transcription start sites exhibit typical eukaryotic features, including +1 nucleosomes, nucleosome-free regions upstream, and phased nucleosome arrays downstream of the TSSs. The global mean nucleosome repeat length varies from 176 bp to 185 bp depending on the developmental stage. Stage specific changes in nucleosome positioning occur locally in intergenic regulatory regions, which are characterized by specific histone modifications and variants. Dynamic nucleosomes correlate with DNA accessibility, gene expression and determine the access to transcription factor binding sites in *Pf*. The highly regular chromatin structure, with stage-specific structural alterations, emphasises the important role of epigenetic mechanisms in regulating the complex life cycles of Pf.

## Introduction

Eukaryotic genomes are organized in the form of a compact nucleoprotein structure, termed chromatin. The basic packaging unit of chromatin is the nucleosome core, which consists of 147 base pairs (bp) of DNA wrapped around an octameric protein core comprising two copies each of the histones H2A, H2B, H3 and H4 (Luger et al. 1997). The nucleosome cores are arranged in continuous arrays separated by short stretches of linker DNA, resembling a beads-on-a-string-like structure (Woodcock et al. 1976). Chromatin is the template for all DNA-dependent processes, and the positioning of nucleosomes on DNA determines the accessibility of DNA sequences to regulatory factors, such as sequence-dependent transcription factors. The positioning, structure and histone modifications of nucleosomes are dynamically changed during signal transduction and developmental processes (Harwood et al. 2019; West et al. 2014; Zhang et al. 2016). The alterations in DNA sequence accessibility that result from these changes establish distinct binding platforms for regulatory factors in different cell types or developmental stages. Minor differences in nucleosome organization can alter the binding behaviour of transcription factors and thus regulate gene activity (Martinez-Campa et al. 2004; J. Li et al. 2006; Längst et al. 1998). Therefore, the analysis and understanding of chromatin organisation and the timing of dynamic changes in nucleosome positioning are crucial to comprehending gene regulatory processes.

In a population of cells, nucleosome positioning can be characterised by three main features: fuzziness, occupancy and position. Nucleosome fuzziness reflects the cell-to-cell variability in nucleosome positioning at genomic sites; higher fuzziness indicates greater variability at individual positions; nucleosome occupancy describes the probability of detecting a nucleosome at a particular genomic position and the position refers to the precise genomic location of the nucleosome dyad. Assessing and quantifying nucleosome occupancy is challenging as methods to map nucleosome positions depend on structural and experimental parameters, such as DNA sequence, non-histone proteins bound to DNA, and nucleosome interactions and stability. These factors impact the quantitative and qualitative isolation or detection of the nucleosomal DNA (Oberbeckmann et al. 2019; Schwartz et al. 2019; Wernig-Zorc et al. 2024; Brogaard et al. 2012). A powerful and widely used method to study these features genome-wide is called micrococcal nuclease sequencing (MNase-seq). MNase-seq uses the property of MNase to preferentially hydrolyse the accessible linker DNA. The histone-bound nucleosomal DNA remaining refractory to MNase cleavage (Yuan et al. 2005; Schwartz et al. 2019), is subsequently sequenced and presents the nucleosome footprint. Several programs exist to analyse MNase-seq data (Shtumpf et al. 2022), such as the commonly used DANPOS toolkit (Chen et al. 2013) or the Nucleosome Dynamics program suite (Buitrago et al. 2019). However, these tools require specific preprocessing of the sequenced DNA fragments and lack the capability to perform comparative analyses across complex experimental designs involving more than two conditions, such as in time series of developmental stages. Recent advancements in bioinformatics pipeline management systems and software containerization enable the development of robust, reproducible, easily deployable and scalable analysis pipelines (Di Tommaso et al. 2017). This approach has been employed to create comprehensive MNase-seq analysis pipelines, such as the nucMACC pipeline, which assesses nucleosome stability and structure (Wernig-Zorc et al. 2024).

The parasite *Plasmodium falciparum* (*Pf*) causes the disease malaria tropica, responsible for 597,000 deaths in 2023 (World Health Organization 2024). It has an intricate life cycle, encompassing various stages in multiple hosts meticulously orchestrated by a complex transcription network (Watzlowik et al. 2021). The just-in-time regulation of transcription requires the precise coordination of chromatin architecture and gene expression throughout the entirety of the developmental process (Watzlowik et al. 2025). Surprisingly, the *Pf* genome encodes only for a reduced set of transcription factors, not matching the need for its complex regulatory network, suggesting that additional regulatory mechanisms must contribute to gene expression control (Bischoff and Vaquero 2010).

The *Pf* chromatin landscape deviates significantly from other eukaryotic systems. The parasite encodes for the most divergent histone sequences in eukaryotes, expresses unique histone variants and notably lacks linker histone H1 (Bischoff and Vaquero 2010). *In vitro* studies have demonstrated that *Pf* nucleosomes possess reduced stability compared to other eukaryotes (Silberhorn et al. 2016). Moreover, chromatin exhibits predominantly euchromatic features, with heterochromatin confined to subtelomeric regions and a few internal islands (S. A. Fraschka et al. 2018). These heterochromatic regions are associated with the regulation of antigenic variation, a mechanism that allows the parasite to evade host immune responses (Flueck et al. 2009). Adding to its unique features, the *Pf* genome is almost devoid of DNA methylation (Ponts et al. 2013) and exhibits an exceptionally high A/T content—averaging 81% across the genome and reaching up to 95% in intergenic regions (Gardner et al. 2002). This high A/T content, combined with the parasite’s “open” chromatin architecture, requires the development of specialized experimental and bioinformatic approaches to analyse the chromatin structure (Bártfai et al. 2010). Micrococcal nuclease (MNase), which has a sequence preference for A/T-rich regions, poses challenges in analysing the *Pf* nucleosome organisation. The parasite’s DNA is more susceptible to endonuclease cleavage, and its nucleosomes are comparatively unstable, leading to potential overdigestion and loss of nucleosomal DNA in MNase-seq experiments (Schwartz et al. 2019; Wernig-Zorc et al. 2024). This has resulted in MNase-seq studies revealing large variations in nucleosome occupancy across the genome with intergenic regions devoid of nucleosomes and irregular nucleosome positioning at intragenic regions (Bunnik et al. 2014; Ponts et al. 2010; Westenberger et al. 2009). These studies proposed that *Pf* chromatin structure is unlike the structure of other eukaryotes. This fact can be explained by the overdigestion of AT-rich DNA with MNase. The currently unique, high-quality MNase-seq dataset systematically spanning the *Pf* intraerythrocytic development cycle (IDC) was generated by Kensche and colleagues using a combination of low MNase concentration and additional Exo III digestion to prevent the overdigestion of AT-rich nucleosomal DNA (Kensche et al. 2016). But still, their analysis, identified only a limited number of well-positioned nucleosomes and failed to detect the regular nucleosome arrays typical of other eukaryotic genomes. These findings, along with others, have contributed to the prevailing view that *Pf* chromatin is atypical, characterized by a loosely organized, highly accessible structure with “fuzzy” nucleosomes and extensive regions of unpackaged DNA (Le Roch et al. 2012; Bunnik et al. 2014; Ponts et al. 2010).

Here we thoroughly re-analyze the stage-specific MNase-seq data from Kensche et al, using our new nucDetective analysis pipeline to improve the understanding of *Pf* chromatin organisation and dynamics along the IDC. nucDetective includes a comprehensive, easy-to-use and state-of-the-art MNase-seq workflow capable of generating high-resolution nucleosome maps starting from raw reads (nucDetective Profiler). Additionally, it offers a multi-condition analysis workflow (nucDetective Inspector), which combines the results from different cellular states to detect the progressive changes in the chromatin landscape. The pipeline presents a universal tool capable of detecting nucleosome dynamics, such as occupancy, fuzziness and position shifts, comparing two or more functional stages. Re-analysis of *Pf* MNase-seq data revealed yet undiscovered chromatin features of the malaria parasite. In contrast to the general assumptions, we show that *Pf* exhibits a chromatin organisation reminiscent of classical eukaryotic chromatin with a well positioned +1 nucleosome, an upstream nucleosome-free region, and phased nucleosome array downstream of the TSS. Improved resolution of nucleosome maps shows changes in chromatin architecture during the IDC, which correlate with DNA accessibility and gene expression, defining actual gene networks being activated or repressed.

## Results

### nucDetective uncovers features of nucleosome organisation and dynamics in *Pf*

We developed nucDetective, an easy-to-use and automated pipeline for analysing MNase-seq datasets enabling the analysis of complex experimental designs, such as time-series experiments. The pipeline was employed to gain deeper insights into chromatin dynamics during the IDC of *Pf*, re-analysing an MNase-seq time-series dataset (Kensche et al. 2016). nucDetective is divided into two consecutive workflows: first, the Profiler, and second, the Inspector.

The nucDetective Profiler workflow begins with raw fastq files, generates high-resolution nucleosome profiles, and identifies nucleosome positions. Alignment and postprocessing steps, such as fragment size selection, are optimised for MNase-seq data, and data quality is controlled at every stage (see Methods for details). Even under challenging conditions, such as the AT-rich *Pf* genome, the results of the Profiler analysis considerably improved the quality of recent nucleosome annotations. A comparison with the previously published MNase-seq analysis clearly shows a gain in structural information providing highly resolved nucleosome positions when using nucDetective Profiler (Figure S1A, S1B and Figure 1A).

**Figure 1.**
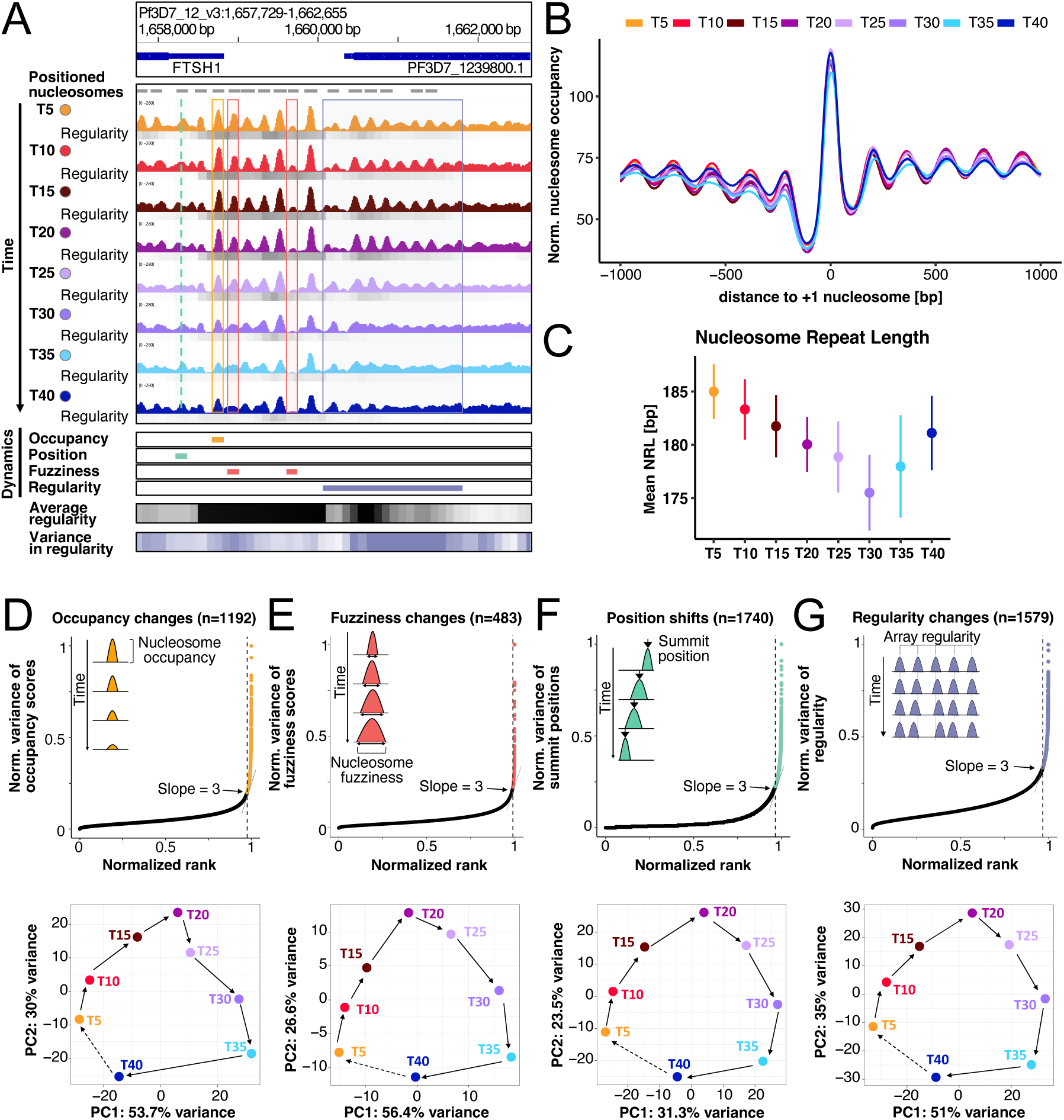
Detection of dynamic nucleosome features in the IDC of *Pf* using the nucDetective pipeline. (A) Genome browser snapshot highlighting the different categories of dynamic nucleosomes. The nucDetective pipeline was used to process MNase-seq data from a time series of the IDC of *Pf* (Kensche et al. 2016). It provides centered nucleosome coverage tracks (T5-T40 colored coverage tracks) and identifies reference nucleosome positions (grey bars). It assesses nucleosome positioning regularity at each timepoint (grey heatmap) and provides an overview of the average regularity (black heatmap) and the variance in regularity (blue heatmap) across all timepoints. Additionally, it detects nucleosomes showing occupancy changes (yellow bar), position shifts (green bar) and fuzziness changes (red bars) as well as regions, where nucleosome positioning regularity has changed (blue bar). Corresponding areas in the nucleosome coverage tracks are marked in this figure with respectively colored lines and rectangles. (B) Nucleosomes upstream and downstream of the TSS in *Pf* are positioned in regular arrays. Average, normalised nucleosome occupancy profiles centered on the +1 nucleosomes are shown. (C) Average genome wide NRL in the IDC of *Pf* changes between 185 bp and 176 bp. The mean NRL with a 95% confidence interval is depicted for each timepoint. The NRL was estimated using the frequencies of same-strand alignment distances through a phasogram. The NRL is determined by the slope of the linear fit to the modes present in the phasogram. (D-G) Dynamic nucleosome calling of (D) occupancy, (E) fuzziness, (F) position and (G) regularity changes. Nucleosomes are sorted by the variance over time of the respective metric. The point where the variance rapidly increases is determined (slope=3) and nucleosomes above this point are considered to reflect changes in occupancy, fuzziness, position or regularity over time. Bottom panels show PCA of the selected nucleosomes. Samples plotted at PC1 and PC2 resemble a cyclic structure as indicated by the arrows reminiscent of the IDC. Percentages in axis labels indicate the proportion of variance explained by each component.

These new *Pf* nucleosome maps reveal a nucleosome organisation at the transcription start site (TSS) reminiscent of the general eukaryotic chromatin structure, featuring a well-positioned +1 nucleosome, an upstream nucleosome-free region (NFR), and a phased nucleosome array downstream of the TSS. Aside from the improved nucleosome resolution, we suggest that the absence of nucleosome arrays downstream of the TSS in previous studies is due to uncertainty in *Pf* TSS annotation and two additional effects. On the one hand, transcription initiation events have been mapped to relatively wide regions in *Pf*, often containing multiple initiation site clusters for a single gene; on the other hand, TSS usage changes during parasite development, leading to divergent TSS annotations (Adjalley et al. 2016; Chappell et al. 2020; Shaw et al. 2021). Aligning nucleosome maps to these variable positions produces inconsistent nucleosome distances, blurring aggregate patterns and obscuring the underlying arrays (Figure S1B). To address this issue, we centered the nucleosome occupancy profile at the positioned +1 nucleosome, using the best positioned nucleosome closest to the assigned TSS within a -100/+300 bp window (Table S1). With this +1 nucleosome annotation, regularly spaced nucleosome arrays downstream of the TSS were detected, revealing a precise nucleosome organisation in *Pf* (Figure S1B). Due to the high resolution maps of nucleosomes we can now observe variations in nucleosome spacing depending on the developmental stage (Figure 1B). For quantifying the average nucleosome repeat length (NRL) at each timepoint, we employed a phasogram-based approach. The largest NRL is noted in the ring stage at T5 (185 bp) and continuously decreases towards trophozoite stage at T30 (176 bp).

The improved nucleosome annotation now facilitates an in-depth analysis of the dynamic changes in the nucleosome landscape during *Pf* IDC (Figure 1A). Genomic regions with regularly spaced nucleosomes, which undergo dramatic structural changes over time, can be clearly identified in the high-resolution data. However, as the data set includes multiple time points, identifying dynamic nucleosomes is not trivial, and MNase-seq optimized tools for analysing such data sets are lacking. To tackle this issue, we developed the nucDetective Inspector workflow.

The Inspector workflow utilises the output of the Profiler workflow to quantify dynamic changes in chromatin structure by analysing multiple nucleosome features: nucleosome occupancy, fuzziness, dyad positions, and the local nucleosome array regularity (Figure 1A, S1C and S1D). To identify dynamic nucleosomes, we employed a similar strategy as that used to define super enhancers or unstable nucleosomes (Whyte et al. 2013; Wernig-Zorc et al. 2024). Nucleosomes are ranked by the variance of the specific feature (occupancy, fuzziness, position, or regularity) across all samples. The variance is normalised to range between 0 and 1 and is plotted against the rank divided by the total number of events (Figure S1C). To geometrically identify dynamic positions where the signal variability increases rapidly, we determined the points on the curve where the slope first exceeds a certain threshold. Here, for the *Pf* analysis, we used a cutoff of 3. For a description of array regularity, the spectral power density of the nucleosome signal at the size of 180 bp was plotted using a rolling window approach (Figure S1D).

The pipeline identified a total of 49.999 reference nucleosome positions from which 1192 nucleosomes exhibiting high variability in occupancy (occupancy changes, Figure 1D), 483 nucleosomes showing high variability in fuzziness (fuzziness changes, Figure 1E), 1740 nucleosomes with pronounced variability in position (position shifts, Figure 1F), and 1579 nucleosomes displaying high variability in local array regularity (regularity changes, Figure 1G). To assess the relevance and information content of the extracted features, a Principal Component Analysis (PCA) was employed. Each set of dynamic nucleosomes exhibits a circle-like data structure in PCA, resembling the progression of *Pf* through every stage of the IDC (Figure 1D-G). In summary, the progressive changes in chromatin structure reflect the continuous developmental process in the life cycle of *Pf*. This finding suggests that changes in chromatin structure are closely associated with the developmental gene expression programme.

### Dynamic nucleosomes reside in regulatory regions and are linked to active promoters

To determine the spatio-temporal distribution of dynamic nucleosomes, we asked whether the various dynamic parameters (position, occupancy, fuzziness) occur at the same or distinct genomic locations. Interestingly, the dynamic parameters show only minor overlaps, indicating that they represent distinct features, potentially associated with specific DNA-dependent processes and chromatin remodeling mechanisms (Figure 2A). At the genomic scale, dynamic nucleosomes are relatively evenly distributed throughout the genome, without apparent feature clustering at specific chromosomal locations (Figure S2A). However, we observe a clear enrichment of dynamic nucleosomes at gene promoters (Figure 2B). This enrichment is particularly prominent for nucleosomes displaying dynamic changes in fuzziness or occupancy. Dynamic alterations in nucleosome fuzziness or occupancy predominantly occur directly upstream of the TSS at the -1 nucleosome regulating the NFR width (Figure 2C). Changes in NFR accessibility or width may be linked to activation or repression of transcription of associated genes (Klein-Brill et al. 2019). In comparison with nucleosomes at the beginning of the gene body, the position of the +1 nucleosome appears to be relatively stable, lacking active position shifts as postulated by the barrier packing model (Mavrich et al. 2008) (Figure 2C). In contrast, nucleosomes exhibiting dynamic changes in array regularity are mainly enriched downstream of the TSS, at the start of the gene body (Figure 2C and S2B), possibly as a consequence of active transcription (Baldi et al. 2018; Singh et al. 2021).

**Figure 2.**
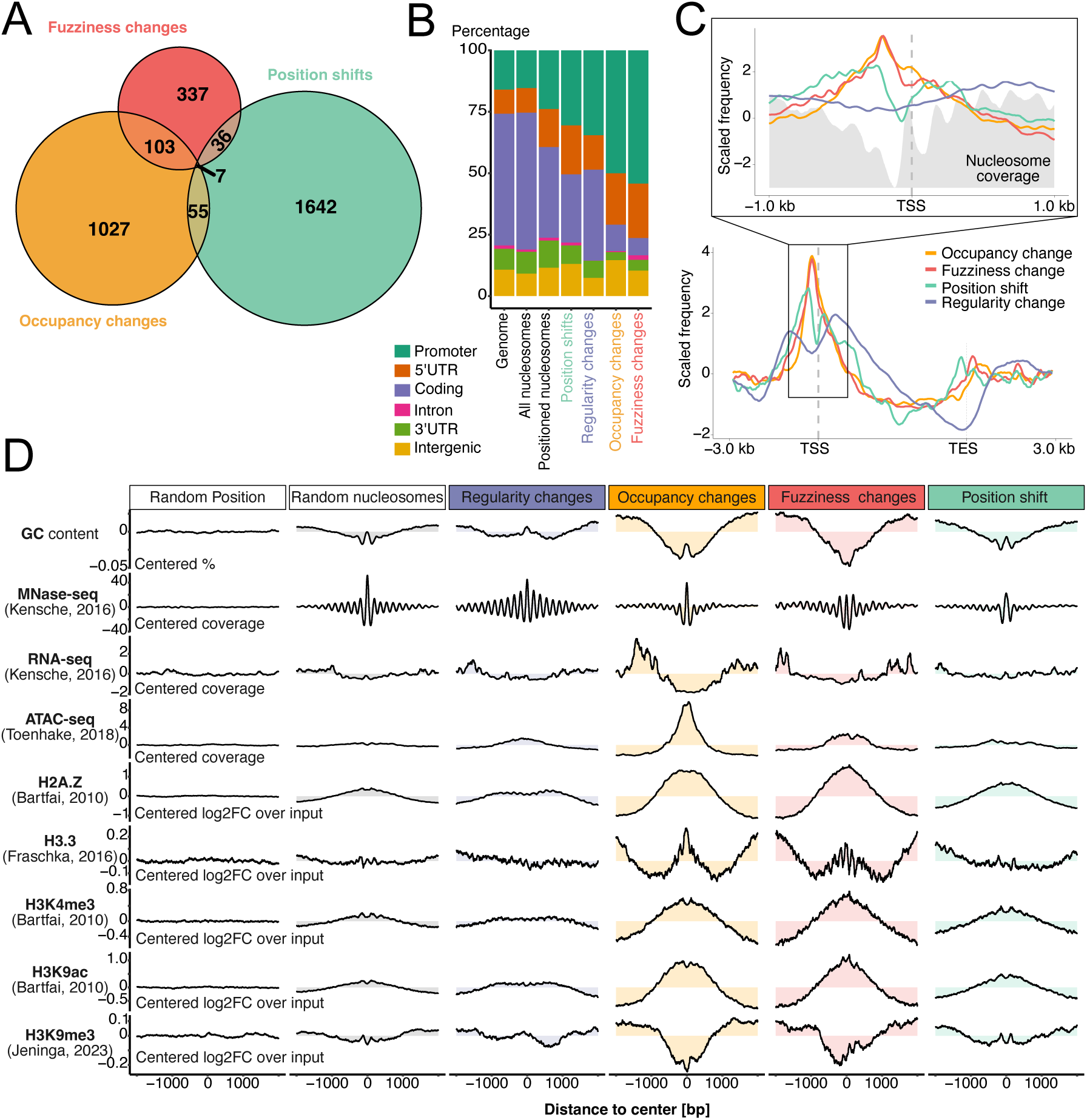
Dynamic nucleosomes reside in regulatory regions and are associated with active promoters. (A) nucDetective characterizes distinct features of dynamic nucleosomes. Euler diagram of dynamic nucleosomes grouped by occupancy (yellow), fuzziness (red) or position changes (green) over time. (B) Dynamic nucleosomes are enriched at the gene promoters. The genome wide distribution of dynamic nucleosomes was assessed at promoter regions (-500 bp to100 bp of the TSS), 5’UTR, coding regions, introns, 3’UTR and intergenic regions using different sets of nucleosomes:: random genomic positions (genome), all called nucleosome positions, well positioned nucleosomes (defined as the 20% with the lowest fuzziness), nucleosomes showing a position shift over time, dynamic nucleosomes showing a change in array regularity, occupancy or fuzziness. (C) Nucleosome occupancy or fuzziness changes and position shifts primarily occur upstream of the TSS at the -1 nucleosome position. The average scaled (z-score normalization) occurrences of nucleosomes exhibiting occupancy change (yellow), fuzziness change (red), position shift (green) and regularity change (blue) is plotted over the scaled gene body. A magnification of the region around the TSS is provided with an unscaled view. For better orientation the inset plot includes the average nucleosome coverage (grey background). (D) Genomic and epigenetic context of dynamic nucleosomes. Average local change compared to genome wide average in GC content, nucleosome coverage, RNA expression, DNA accessibility measured by ATAC-seq, histone variants H2A.Z and H3.3 and the histone modifications H3K4me3, H3K9ac and H3K9me3 are plotted at random genome positions, random nucleosome dyads and dyads of dynamic nucleosome categories. Shaded areas illustrate the deviation to the genome wide average.

Next, the dynamic nucleosome categories were compared to previously published studies on histone variants (S. A.-K. Fraschka et al. 2016; Bártfai et al. 2010), histone modifications (Bártfai et al. 2010; Jeninga et al. 2023), DNA-accessibility (Toenhake et al. 2018) and transcription (Kensche et al. 2016). These and other studies suggested that the histone variant H2A.Z is a marker for regulatory regions in *Plasmodium*, guiding chromatin modifying and transcription initiating complexes (Bártfai et al. 2010). H2A.Z occupancy remains constant throughout the erythrocytic lifecycle, whereas the H3K4me3 and H3K9ac histone marks associated with H2A.Z are stage-specific and correlate with the regulation of the developmental cycle (Bártfai et al. 2010). The H3.3 variant binding sites in *Pf* are suggested to depend on the GC content of DNA, marking coding and subtelomeric repetitive regions, irrespective of transcriptional activity (S. A.-K. Fraschka et al. 2016). Our analysis shows that dynamic nucleosomes, changing occupancy and fuzziness, preferentially occur in the H2A.Z/H3K4me3/H3K9ac marked regions and are also linked to regions containing the histone variant H3.3 (Figure 2D).

Heterochromatin in *Plasmodium falciparum* is characterised by the presence of H3K9me3 and heterochromatin protein 1 (HP1). It is observed in subtelomeric regions and small internal regions where it is involved in silencing virulence factors such as multi-gene surface antigens, while also playing a role in life cycle stage transitions (S. A. Fraschka et al. 2018; Flueck et al. 2009). Heterochromatin domains, as indicated by H3K9me3 ChIP, are characterised by depleted nucleosome occupancy and fuzziness changes, indicating a stable chromatin organisation (Figure 2D). This underscores the tight epigenetic control of these essential regions involved in parasite adaptation and survival. While there is a strong association between fuzziness and occupancy dynamics with the histone variants and the H3K4me3/H3K9ac marks, the data can be further subdivided according to the ATAC-seq pattern. We observed a strong ATAC peak at nucleosome positions undergoing occupancy changes; however, only a few ATAC sites coincided with changes in nucleosome fuzziness, and even fewer with position shifts (Figure 2D). In summary, nucleosome dynamics is intimately connected with specific histone modifications and variants at active genomic sites, primarily located at gene promoters, emphasising their essential role in regulating gene expression.

### Dynamic nucleosomes (anti-)correlate with DNA accessibility but exhibit distinct features

Next, we examined the dynamics of nucleosomes in accessible chromatin regions, defined by ATAC-seq positive domains (Toenhake et al. 2018). The Profiler workflow annotated a total of 5300 nucleosomes in these open chromatin domains (Figure 3A). While most of these nucleosomes remained stable over time (n=4129) a significant subset (n=1171, p < 0.0001, permutation test) of nucleosome positions exhibited a dynamic behaviour (Figure 3A). As indicated above (Figure 2D), open chromatin regions are predominantly associated with changes in nucleosome occupancy, accounting for approximately 58 % of all nucleosomes in this class (n=689, p < 0.0001, permutation test) (Figure 3A). Analysis of the association between chromatin accessibility and nucleosome occupancy dynamics revealed a negative relationship (median Pearson correlation of ρ = -0.635) (Figure 3B-C), indicating that a decrease in nucleosome occupancy accompanies chromatin opening. Notably, we observed that the eviction of a single nucleosome is sufficient to open broader regions (Figure 3C). Similarly, but to a lesser extent, we observed a positive correlation with nucleosome fuzziness (median Pearson correlation of ρ = 0.485) (Figure 3A and 3B). However, nucleosome shifts are rarely present in open regions (15%, n=261) and do not correlate with chromatin accessibility (Figure 3A-C). As ATAC-seq is not suitable for resolving nucleosome shifts, not all chromatin features can be detected by this method. MNase-seq analysis by the nucDetective pipeline achieves a more comprehensive and better resolved view of chromatin dynamics.

**Figure 3.**
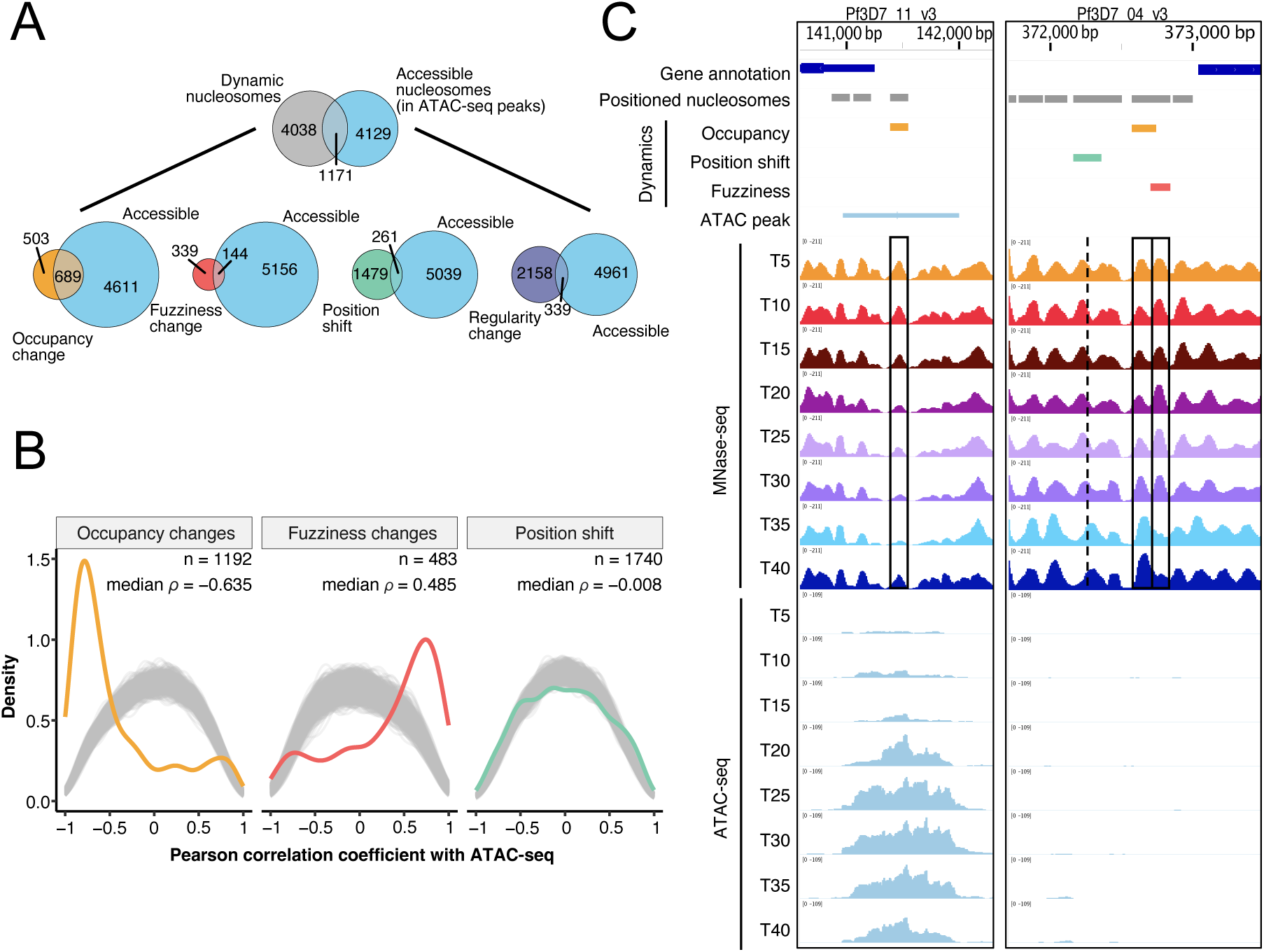
Dynamic nucleosomes (anti-)correlate with DNA accessibility yet show distinct features. (A) Mainly nucleosome occupancy and fuzziness changes coincide with open chromatin regions. Venn diagrams illustrating the overlap of nucleosomes in open chromatin regions derived from ATAC-seq in at least one timepoint (lightblue) and dynamic nucleosomes (grey). Below the overlap with individual nucleosome features are shown. (B) Loss of nucleosome occupancy and increase in fuzziness are correlated with DNA accessibility changes. Linear correlations were computed between the accessibility score derived from the ATAC-seq fragment coverage at each nucleosome position and the corresponding occupancy, fuzziness and summit position (shift) at every timepoint. The density plot illustrates the Pearson correlation coefficients of dynamic nucleosomes, alongside the density of Pearson correlations for randomly selected, equally sized sets of nucleosomes (grey, 1000 iterations). The median Pearson correlation coefficient *ρ* and the number of nucleosomes (n) are indicated. (C) Genome browser snap shot showing representative examples of the anti-correlation between nucleosome occupancy and DNA accessibility (left) and a region showing nucleosome dynamics but no changes in DNA accessibility (right). Black boxes and lines are provided for easier visual identification of dynamic nucleosomes.

### Transcription factor binding motifs are associated with distinct nucleosome occupancy kinetics

Nucleosomal stability and positioning regulate the availability of specific DNA elements for regulatory proteins (Zhu et al. 2018). Modulating nucleosome positioning at specific sites affects transcription factor binding and ultimately controls gene expression programs. Therefore, we examined the kinetics of changes in nucleosome occupancy in greater detail. Clustering analysis revealed groups of nucleosomes that open and close in a concerted manner at different time points of the erythrocytic life cycle (Figure 4A). Cluster 1 comprises genomic sites with low nucleosome occupancy during the first 20 hours of the erythrocytic life cycle, thereby allowing regulatory proteins access to the underlying DNA. Starting at 25 hours, nucleosome deposition occurs at these sites, thus restricting factor access to these DNA sequences. To uncover recurring sequence elements in regions characterized by similar nucleosome kinetics, we performed de novo motif analysis (Figure S3), and identified motifs were compared to known ApiAP2 transcription factor motifs (Campbell et al. 2010) (Figure S3, last 3 columns). ApiAP2 transcription factors are, with 27 known members, the largest family of transcription factors in *Pf*. These are involved in the regulation of IDC progression and differentiation (Balaji et al. 2005; Campbell et al. 2010). Several overrepresented motifs displayed high similarities to the binding motifs of the ApiAP2 transcription factor family, such as AP2-FG, AP2-O4, AP2-G5 and AP2-I (Figure 4B). Remarkably, the time point of nucleosome eviction in the nucleosome group (cluster 5) associated with the AP2-I transcription factor motif correlates with the peak of AP2-I expression and its putative role in erythrocyte invasion (Santos et al. 2017). The analysis also revealed novel sequence motifs (Figure 4), suggesting the existence of other sequence specific DNA binding factors in *Pf*, binding to regulatory elements and playing a role in the intricate regulation of the life cycle of *Pf*.

**Figure 4.**
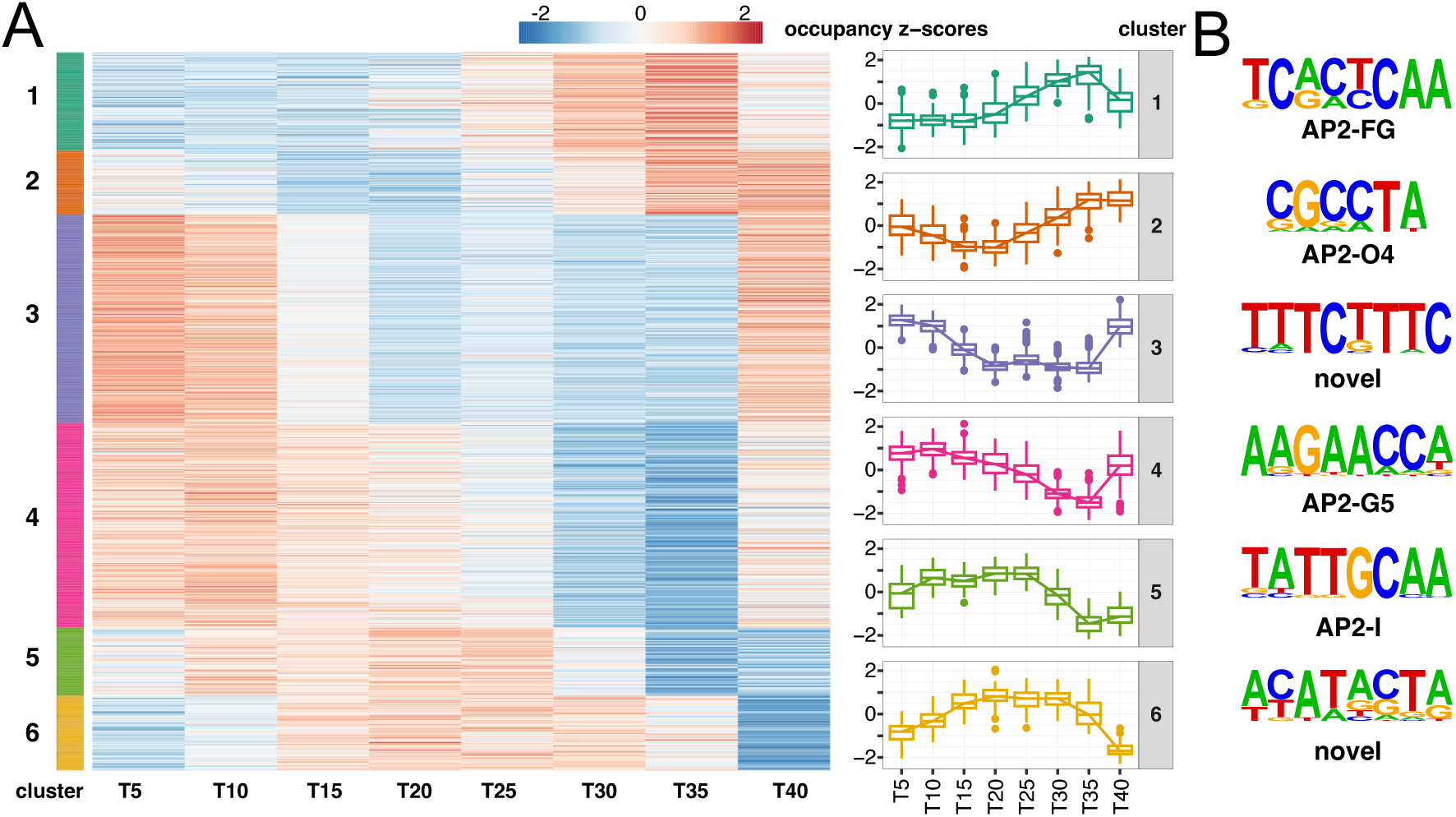
Transcription factor binding motifs are associated with distinct nucleosome occupancy kinetics. (A) Nucleosome occupancy dynamics can be grouped by distinct kinetics in the lifecycle of *Pf.* Self-organizing map clustering of nucleosome occupancy changes resulted in 6 distinct clusters. Heatmap (left) showing z-score scaled occupancy scores across all time points ordered by cluster assignment as indicated by the color bar on the left side. Boxplots (right) depicting the z-score scaled occupancy changes over time of the individual clusters as indicated on the right side. (B) Distinct DNA motifs are associated with nucleosome occupancy kinetics. The top *de novo*– derived motif for each cluster is shown. The transcription factor names of the best-matching known DNA binding motifs derived from (Campbell et al. 2010) are shown. Motifs where we did not find a corresponding known DNA binding motif (similarity score < 0.6) are marked as novel. Additional enriched motifs along with the significance of motif enrichment and the fraction of motifs at the respective nucleosome positions are shown in Figure S3

### Nucleosome dynamics in promoter regions are associated with gene expression changes

The accumulation of dynamic nucleosomes in promoter regions (Figure 2B) suggests a role for nucleosome positioning in the regulation of gene expression. The high-resolution maps of nucleosome positions obtained with the nucDetective pipeline allow for the first time, a detailed analysis of the structural changes at *Pf* promoter regions. Selecting genes that are activated upon entry into the trophozoite stage at T25 shows a concomitant opening of the promoter region and the formation of a nucleosome-depleted region upstream of the +1 nucleosome (Figure 5A and 5B). Active transcription also results in a loss of regularly positioned nucleosomes in the gene body (Figure 5A-B, S4A). However, by T40, the chromatin structure of these genes is completely reverted, closing the promoter and re-establishing the regular nucleosome array in the gene body. This can be observed even though the transcript abundance remains high, as shown by the RNA-seq data (Figure 5A). It must be noted that the RNA-seq data provides steady-state transcript levels, not allowing the conclusion that chromatin closure is occurring during active transcription. Consistently, nascent RNAs detected by GRO-Seq reveal transcriptional repression of this gene set in the late schizont stage, correlating with promoter closure (Lu et al. 2017) (Figure S4B). Furthermore, classifying genes according to their nascent transcript profiles into four groups reveals characteristic chromatin structures at the gene promoters, temporally correlating with ongoing transcription (Figure S4C). Interestingly, as most genes are repressed in the early ring and late schizont stage, we observe a high similarity in the promoter chromatin structure for T5 and T40, except for those genes which are expressed late (Figure S4C, cluster A8).

**Figure 5.**
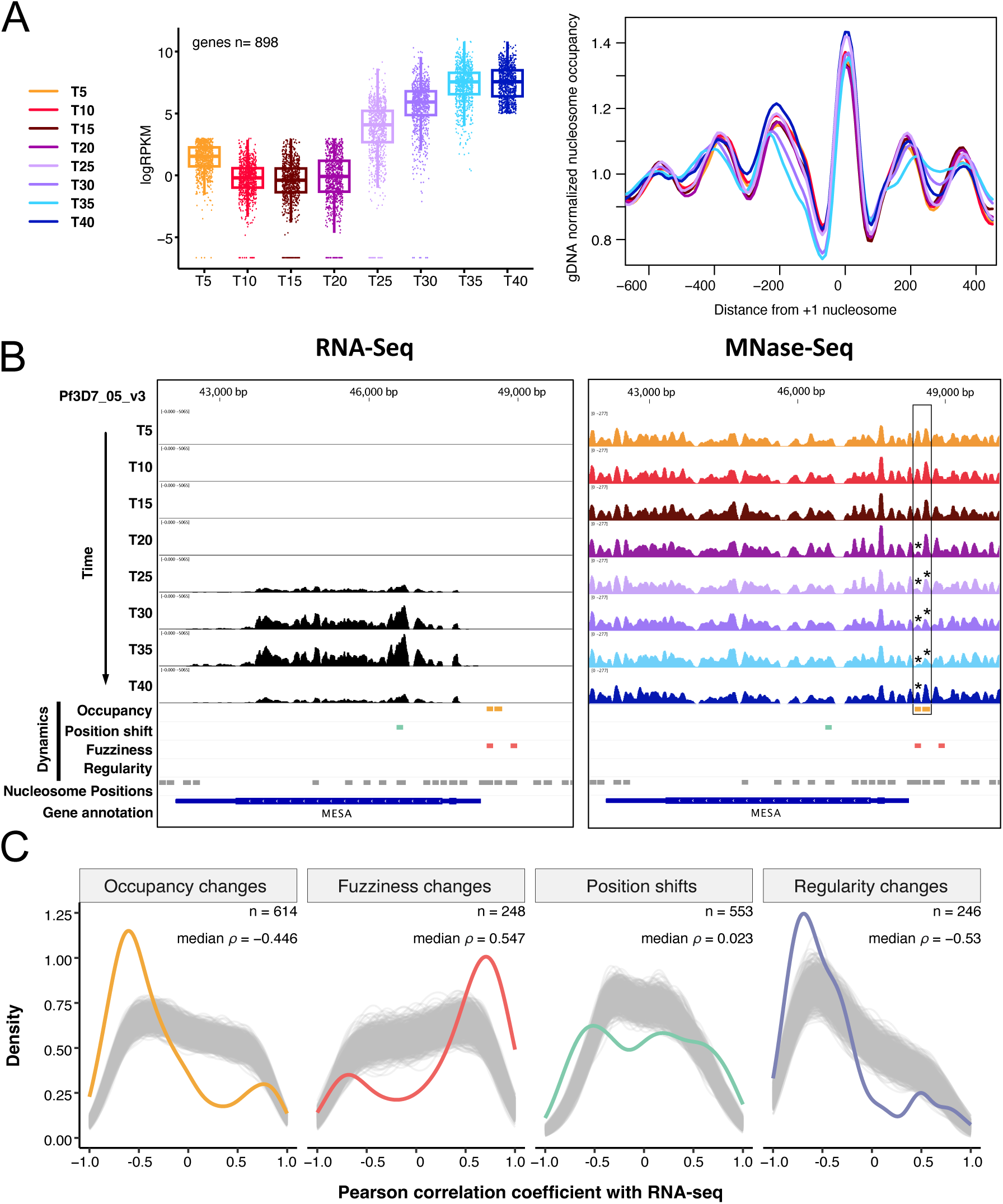
Nucleosome dynamics in promoter regions are associated with gene expression changes. (A) The opening of the promoter region correlates with the initiation of transcription. Genes with a low transcript abundance at the beginning and a high abundance at the end of the IDC were selected (n = 898, left). Average nucleosome occupancy profiles centered at the +1 nucleosomes show an opening of promoter region at T30 and T40 resulting in a nucleosome-depleted region upstream of the TSS (right panel). Nucleosome occupancy profiles were first scaled by the underlying profile of MNase digested gDNA and then the scaled coverage profile at each time point was divided by its region median coverage value. (B) Genome browser snapshot illustrating the correlation between nucleosome dynamics (right) and gene expression (left). Gene expression becomes detectable simultaneously with nucleosome eviction upstream of the TSS (black box, yellow bars). Stars mark the changes in nucleosome occupancy. (C) Nucleosome eviction, loss of position regularity and an increase in fuzziness in promoter regions (-500 to +100bp from TSS) correlate with transcriptional activity. Linear correlations were computed for each dynamic nucleosome in the promoter region and the corresponding gene expression. The density plot illustrates Pearson correlation coefficients for dynamic nucleosomes, alongside the density of Pearson correlations for randomly selected, equally sized sets of nucleosomes in promoter areas (grey, 1000 iterations). The median Pearson correlation coefficient *ρ* and the number of nucleosomes (n) are indicated.

The data shows globally that transcriptional effects are associated with dynamic nucleosomes in promoter regions (-500 to + 100 bp from TSS), where nucleosome eviction (loss of occupancy, median Pearson correlation ρ = -0.446) and loss of positioning (increased fuzziness and loss of regularity, median Pearson correlations of ρ = 0.547 and ρ = -0.53 respectively) correlate with transcriptional activity (Figure 5C). A similar trend is observed for all nucleosomes in coding regions, where occupancy, fuzziness and regularity (anti-)correlate with gene expression (Figure S4A).

## Discussion

We developed an optimised MNase-seq analysis pipeline called nucDetective, which is designed to annotate nucleosome positions at high resolution (Profiler workflow) and to identify dynamic changes in nucleosome positioning (Inspector workflow). The Profiler workflow commences with raw sequencing files, automates the process to produce high-resolution nucleosome profiles, and incorporates necessary quality checks (QC), making it a universal tool for any MNase-seq dataset. This is achieved using MNase-seq optimized alignment settings, and proper selection of the fragment sizes corresponding to mono-nucleosomal DNA to increase resolution. Implementing these features into a seamless workflow resulted in a clearer definition of nucleosome positions and detection of well-positioned nucleosomes. In this study, we demonstrate that the Profiler workflow effectively analyses challenging MNase-seq data in *Pf*.

The consecutive Inspector workflow allows for a comprehensive comparison of nucleosome features for experimental designs that exceed a simple two-condition comparison, setting it apart from other MNase-seq analysis tools. The method is versatile and can be used in various experimental settings, such as cell differentiation, knock-down approaches, and time-series treatment experiments, or even for comparing multiple functional states, like the IDC of *Pf* presented here. Simultaneously assessing changes in nucleosome occupancy, position, fuzziness, and array regularity enables an in-depth analysis of chromatin structure dynamics. This provides a higher resolution and more insights into nucleosome features than is possible using ATAC-seq. The nucDetective pipeline is user-friendly, allowing experimental scientists to utilise a state-of-the-art pipeline with relevant QC metrics and implement required software packages without complex installation. Its implementation as an open-source nextflow pipeline allows for modular extension and adaptation to emerging demands in the future (Di Tommaso et al. 2017).

Here, we re-analysed the MNase-seq dataset of Kensche and colleagues to investigate the poorly understood chromatin architecture of *Pf*. The parasite exhibits a complex life cycle in two hosts, revealing dramatic changes in its gene expression programme. A scarcity of transcription factors suggests a large contribution of epigenetic mechanisms to transcriptional regulation (Templeton et al. 2004). Indeed, specialised chromatin remodelling enzymes organise chromatin architecture temporally and structurally, shaping the interaction landscape for transcription factors (Watzlowik et al. 2021; 2025). To better understand these epigenetic processes, we provide a comprehensive analysis of the nucleosome landscape and its dynamics in *Pf* in unprecedented detail. Our results demonstrate that *Pf* exhibits a typical eukaryotic chromatin structure, including well-defined nucleosome positioning at the TSS and regularly spaced nucleosome arrays (Schones et al. 2008; Yuan et al. 2005), which has not been clear in the field up to now. Furthermore, we show temporal dynamics of nucleosomes and link these to chromatin accessibility and gene expression patterns. Our analysis also identifies putative cis-regulatory elements that may operate in a coordinated manner during transcriptional regulation. Together, these findings advance our understanding of chromatin-based gene regulation in *Pf* and challenge prevailing views about its chromatin architecture.

Studies on *Pf* have demonstrated that it has a general transcription machinery akin to that of other eukaryotes, which implies the existence of a classical promoter organisation (Bischoff and Vaquero 2010). However, using the annotated TSS or the translational start site (ATG) as reference points to visualise nucleosome positions did not reveal a eukaryotic-like chromatin architecture (Kensche et al. 2016). However, the TSS in *Pf* is not well-defined, occurring within a relatively broad window rather than at a single, fixed location (Adjalley et al. 2016). Furthermore, the TSS varies according to the developmental stage of the parasite, often using multiple TSS windows and multiple promoters for the same gene (Adjalley et al. 2016), adding additional complexity to TSS annotation. The ATG codon, on the other hand, is also suboptimal as a reference point due to its variable distance from the TSS and its independence from transcription initiation and the associated chromatin structure. By focusing on the nucleosome closest to the annotated TSS (the +1 nucleosome) as a reference point in this study, we were able to obtain a clear nucleosome positioning pattern at promoters, demonstrating the important role of the +1 nucleosome in determining the transcriptionally competent promoter structure. Accordingly, we show that the chromatin structure of the *Pf* promoter indeed resembles a typical eukaryotic promoter. The improved resolution of nucleosome positions also enables us to calculate the average nucleosome repeat length (NRL) throughout the parasite’s life cycle. Early studies on *Pf* chromatin structure suggested very short NRLs of 150 to 155 bp at specific genes, which was confirmed using the fragment length of isolated di-nucleosomes in MNase-seq experiments (Silberhorn et al. 2016; Lanzer et al. 1994; Horrocks et al. 2002). However, employing the nucDetective pipeline on the Kensche dataset allows to assess the average genome wide NRL, which range from 176 to 185 bp and align with the typical sizes of eukaryotic NRL‘s (Compton et al. 1976; Prunell and Kornberg 1982). Our result are also consistent with previous findings that cells with high transcriptional activity exhibit a reduction in nucleosome-to-nucleosome distance (Baldi et al. 2018). We observed the shortest NRL in the trophozoite stage of *Pf* which coincides with elevated transcriptional activity and significant opening of chromatin (Bunnik et al. 2014; Ay et al. 2014; Ponts et al. 2010; Lu et al. 2017; Kensche et al. 2016). Conversely, the longest NRL is observed during the ring stage, shortly after erythrocyte invasion, when transcriptional activity is very minimal (Kensche et al. 2016; Lu et al. 2017). This suggests a potential link between nucleosome spacing and transcriptional activity in *Pf*.

We demonstrate that nucleosome dynamics occur predominantly in the promoter region of genes in *Pf*, and we can trace the alterations to specific nucleosomes and genes. We find that the specific regulation of individual nucleosomes directly correlates with changes in accessibility and downstream transcription, thereby enhancing the resolution of biologically relevant regions of chromatin as compared to other methods, such as ATAC-seq. Furthermore, the analysis of concerted nucleosome dynamics has facilitated the identification of novel sequence motifs that become accessible, revealing potential regulatory elements and suggesting the existence of yet undiscovered DNA sequence specific factors. Our discovery supports previous studies indicating that there are undiscovered Plasmodium-specific transcription factors (Bischoff and Vaquero 2010; Militello et al. 2004). Here we demonstrate that our high-resolution chromatin structure analysis provides new insights into the complex regulatory network. Moreover, apart from the information revealed by ATAC-seq data, which monitors chromatin accessibility, we show that nucleosome position shifts do not result in chromatin opening and are not detected by ATAC-seq. The function of these nucleosome position switches still needs to be elucidated. The nucDetective pipeline highlights the unique ability of MNase-seq to extract relevant features that cannot be inferred from other methods used to probe chromatin structure. Investigating the processes associated with chromatin dynamics that do not lead to increased accessibility may present an interesting area of research in the future.

## Materials and Methods

### The nucDetective pipeline

All the steps from fastq files to detection of dynamic nucleosome features across multiple conditions were integrated and automated within the nucDetective pipeline, which is available on GitHub (https://github.com/uschwartz/nucDetective.git). The version of the nucDetective pipeline used in this study was v1.1. The nucDetective pipeline is executed using nextflow workflow management system and runs the software within stable Docker containers, ensuring a reproducible, scalable and portable analysis workflow (Di Tommaso et al. 2017). The nucDetecitve pipeline is split into: 1) the Profiler workflow, comprising mapping, pre-/post-processing, QCs, NRL analysis and nucleosome profiling, and 2) the consecutive Inspector workflow, comprising nucleosome profile normalization, exploratory data analysis, nucleosome position annotation and detection of dynamic nucleosome features.

### The Profiler workflow

The input of the workflow are raw paired-end sequencing data in the form of fastq files. First sequencing and read quality is checked using FastQC (Andrews 2010) and adaptor sequences or low quality bases are trimmed using Trim Galore with the parameter: --stringency 2 and -q 10 (Krueger 2023). Reads are mapped against the indexed reference genome (here Pfalciparium3D7 release 57 from PlasmoDB) using bowtie2 with following settings: --very-sensitive, --no-discordant, --no-mix, --dovetail (Langmead and Salzberg 2012). Aligned fragments are filtered by MAPQ scores of at least 20 and reads mapped in proper pair using samtools (H. Li et al. 2009). Quality of the aligned sequence data, such as the fragment length distribution, is controlled using qualimap (García-Alcalde et al. 2012). The fragments are further filtered to mono-nucleosome sized fragments (here we used 75 – 175 bp) and optionally fragments mapping to blacklisted elements are removed (here the mitochondrial and apicoplast chromosome) using the alignmentSieve function of deepTools package (Ramírez et al. 2016). Nucleosome fragment coverage profiles are normalized by the supplied mappable genome size (here 23292622 bp) and generated using the dpos function of the DANPOS2 package with following options: -m 1, --extend 70, -u 0, -z 20, -e 1, --distance 75 –width 10 (Chen et al. 2013). Nucleosome repeat lengths (NRLs) are calculated based on a phasogram approach described in (Valouev et al. 2011) and implemented in the swissknife (v0.40) package. Frequencies of distances between 5’ ends of reads in the same orientation are calculated and linear regression between peak maxima allows estimation of the average phase between nucleosomes.

### The Inspector workflow

The nucleosome fragment coverage profiles in the form of wig files, which are the result of the Profiler workflow, are used in the consecutive Inspector workflow as input. First the nucleosome profiles are quantile normalized to a selected reference profile (here T20) using the wiq function of the DANPOS2 package (Chen et al. 2013). Optionally, if a TSS annotation is provided (here we used our +1 nucleosome centered TSS annotation, Table1), a TSS plot of the normalized profiles can be generated using the computeMatrix and plotProfile functions of the deepTools package (Ramírez et al. 2016). Next, nucleosome positions of each sample are called using the dpos function of DANPOS2 package with following settings: -z 20, -e 1, -- width 10, --height 25. The nucleosome annotation result of DANPOS2 is converted to bed file format and the best 20% positioned nucleosomes in each sample are filtered based on the provided fuzziness score. Next, the selected nucleosome positions of each sample are stepwise merged into a common reference nucleosome map. Nucleosome positions that overlap by at least 100 bp are stitched together. Nucleosome occupancy of the reference nucleosome map is assessed in each sample using the deeptools multiBigwigSummary function on the normalized nucleosome profiles. The resulting nucleosome occupancy table is than used for exploratory data analysis, such as PCA and correlation clustering.

### Nucleosome fuzziness, occupancy, shift and regularity dynamics

Nucleosome fuzziness scores as reported from DANPOS2 and nucleosome occupancy scores assessed with multiBigwigSummary (see above) are taken for subsequent analysis. Nucleosome occupancy scores are further rlog transformed to minimize differences between samples and stabilize the variance (Love et al. 2014). To detect nucleosome shifts the nucleosome profiles are loaded for each nucleosome position on the nucleosome reference map and a locally weighted scatterplot smoothing (LOESS) is applied using an alpha parameter of 0.6. The summit of the smoothed curve is assessed as nucleosome dyad position in each sample and used for further analysis.

To quantify nucleosome regularity, spectral power density (PSD) analysis was applied to nucleosome occupancy profiles derived from normalized bigWig coverage files. Genomic coverage was extracted and processed in a rolling window manner (width: 1025 bp; step size: 100 bp), and power spectra were estimated using the smoothed periodogram function with a spectral smoothing span of 3 and a padding factor of 1. Spectral components were computed for each window, and the nucleosome repeat length (NRL) was derived as the inverse of the frequency. For each window, the spectral power at the frequency nearest to the expected average NRL (180 bp) was extracted, yielding a genome-wide track of regularity scores. These scores were log-transformed and used to compute variance and mean tracks across all samples. Regularity scores were assigned to individual nucleosomes by averaging PSD values over centered bins spanning 800 bp around each reference nucleosome dyad.

To identify the most dynamic nucleosomes in regularity, fuzziness, occupancy or position (shift) the respective scores are normalized to a deviation from the highest to lowest value of 1 and plotted against their ranks, which are normalized by the total number of ranks. A LOESS smoothing was applied and the first derivate of the LOESS fit was calculated to deduce the slope of the curve. The most dynamic nucleosomes are determined as the highest scores after the slope of the curve exceeds 3.

### +1 nucleosome annotation

To get +1 nucleosome annotation, mono-nucleosome sized fragments of all timepoints were merged and used as input for Nucleosome Dynamics program suite with PlasmoDB57 as reference annotation (Buitrago et al. 2019). The 3’-end of the resulting gff from txstart was used as +1 nucleosome dyad annotation for all timepoints. Nucleosome coverage maps are aligned to regions +/- 1000 bp around the +1 dyad annotation and the average coverage is plotted.

## Downstream Analysis

PCA was performed using nucleosome features (occupancy, fuzziness, position, regularity) of each timepoint on nucleosomes with high variance of the respective features (e.g. dynamic nucleosomes; results of nucDetective Inspector).

Overlaps of nucleosome labels were visualised using the eulerR package (Larsson 2024). The genome wide distribution of dynamic nucleosomes was assessed using the ChIPseeker package with PlasmoDB57 annotation (Yu et al. 2015). Promoter region was defined as -500 to 100 bp around the TSS. The frequency profile of dynamic nucleosomes at the TSS and in gene bodies uses PlasmoDB56 annotation and deeptools for meta- and composite plots (Ramírez et al. 2016). The profiles are scaled and smoothed by a rolling average with window length of 25 bp and 5 bp for gene bodies and TSS area plots respectively.

### Association with histone marks

For comparability samples from ring stages or 20 hours past infection were used. Datasets for H3K9me3 (GSE202214) (Jeninga et al. 2023), H3K9ac, H3K9me3 and H2A.Z (GSE23787) (Bártfai et al. 2010) and H3.3 (GSE80466) (S. A.-K. Fraschka et al. 2016) were analysed as follows: Reads were trimmed with trimmomatic v0.39 using the following options: ILLUMINACLIP:<AdapterSequences>:2:30:10 MAXINFO:30:0.2 MINLEN:35. Alignment was done using bowtie2 v2.5.1 (Langmead and Salzberg 2012) with “--very-sensitive “, “--no-mixed “ and “--no-unal “ options. Further processing was done with deeptools v3.5.1 (Ramírez et al. 2016). First bamCoverage with options “-bs 10 --normalizeUsing RPKM “ was used to get coverage bigwigs and subsequently bamCoverage -b1 <ChIP.bw> -b2 <Input.bw> -bs 10 – operation log2 was used to get log2(ChIP/Input) bigwigs.

ATAC-seq data was obtained from GEO (GSE104075) (Toenhake et al. 2018) as bedgraph files, which were converted to bigwigs using the ucsc bedgraphtobigwig tool. RNA-seq data (GSE66185) (Kensche et al. 2016) were downloaded and adjusted to PlasmoDB57 annotation with 10 bp stepsize by a custom R script. bedtools v2.30.0 “makewindows” and “nuc” functions were used to get the GC content of the *Pf* genome in 150 bp windows (Quinlan and Hall 2010). The average, centred log2FC or coverage of histone marks, RNA-, MNase- and ATAC-seq around nucleosome positions is plotted.

### Overlaps and Correlation of Dynamic Nucleosomes with ATAC Peaks

ATAC peaks are obtained from GEO (GSE104075) (Toenhake et al. 2018) Nucleosomes that overlap by at least 100 bp were assigned to the corresponding ATAC peak. Overlaps are visualised using the eulerR package (Larsson 2024).

### Correlation of nucleosome features with accessibility

Chromatin accessibility at the eight developmental timepoints was quantified by averaging ATAC-seq signal intensity across each reference nucleosome position. Fuzziness scores, occupancy, and positional shift data for the reference nucleosomes were obtained from the nucDetective Inspector workflow. For each nucleosome, Pearson correlation coefficients between ATAC measured accessibility and individual nucleosome features were calculated across all timepoints. To assess the statistical significance of these correlations a permutation-based approach was applied. Null distributions were generated by randomly sampling nucleosomes from the genome 999 times per feature. Correlation values from the set of highly dynamic nucleosomes were compared to these simulated backgrounds in the promoter regions.

### Clustering of nucleosome occupancy dynamics

Self-organising maps (SOM) were used to group changes in nucleosome occupancy according to the time at which they occurred (Wehrens and Buydens 2007). A hexagonal grid was initialised to the size of 13 x 13. The learning rate was set to α = (0.05,0.01) and the number of iterations was set to 500. Hierarchical clustering was applied to the build SOM to obtain six cluster using the agglomeration method “ward.D” in the R function hclust.

### Motif analysis of nucleosome occupancy cluster

The function findMotifsGenome of the HOMER suit was applied to the nucleosome positions of each cluster to find enriched sequence motifs (Heinz et al. 2010). As background the 20% best positioned nucleosome reference map was provided. Identified *de novo* motifs were compared to the set of known motifs for transcription factors of the ApiAP2 protein family from *Pf* (Campbell et al. 2010). For each cluster the three best ranked (according to their p-value) *de novo* motifs were reported. Furthermore, for each of these top ranked motifs matches with known motifs are shown using a similarity score cutoff of 0.6. If the similarity score was below this threshold, only the top match was included.

### Correlation of nucleosome features with gene expression

Nucleosomes from the reference nucleosome map (result of nucDetective Inspector) were assigned to genes according to their respective promoters (500 bp upstream to 100 bp downstream from TSS) or coding regions. Gene expression data (GSE66185) in the form of rescaled RPKM values were adopted from Kensche et al. (Kensche et al. 2016). The gene expression data and nucleosome features (fuzziness, occupancy and shifts) were aggregated and the Pearson correlation was calculated for each nucleosome and feature across all developmental time points. The statistical significance of these correlations was assessed utilizing a permutation-based approach, randomly sampling nucleosomes from the respective genomic region (promoter or coding regions) 999 times per feature to create a null distribution. The correlation values from dynamic nucleosomes were compared to these simulated distributions in the promoter region and to all nucleosomes in the coding regions.

### Public datasets used in this study

*Pf* MNase-seq data and RNA-seq data analysed in this paper are available at GEO with accession GSE66185. Datasets for H3K9me3 (GSE202214), H3K9ac, H3K9me3 and H2A.Z (GSE23787), H3.3 (GSE80466) and ATAC-seq (GSE104075) were obtained from GEO.

## Acknowledgments

We thank Richard Bartfai and Manuel Llinas for fruitful discussions and comments on this manuscript. The work was funded by the Detusche Forschungsgemeinschaft (DFG) with the project number 534335380.

## Author contributions

Conceptualization: US, SH, GL, MTW

Methodology: US, SH, LS, VMR

Formal analysis: US, SH, LS, VMR

Investigation: US, SH, LS, VMR

Visualization: US, SH, LS

Software: US, SH, LS, VMR

Supervision: US, GL

Funding Acquisition: GL

Resources: GL

Writing—original draft: US, SH, GL

Writing—review & editing: US, SH, GL

## Data and materials availability

All data needed to evaluate the conclusions in the paper are present in the paper and/or the supplementary Materials. The nucDetective pipeline v1.1 is available on GitHub with the repository <u>uschwartz/nucDetective</u>. Additional scripts to replicate the analysis presented in this paper are found at <u>SimHolz/Holzinger_et_al_2025</u>.

The results of the nucDetective *Pf* analysis including nucleosome profiles, nucleosome reference positions and dynamic nucleosomes, as well as the used *Pf* annotation are deposited on https://zenodo.org/ with the DOI: 10.5281/zenodo.16779899.

