## Supplementary Material for "Deciphering chromatin architecture and dynamics in *Plasmodium falciparum* using the nucDetective pipeline"

Simon Holzinger *et al.*

**This PDF file includes:**

Figures S1 to S4

**Other Supplementary Materials for this manuscript include the following:**
Table S1

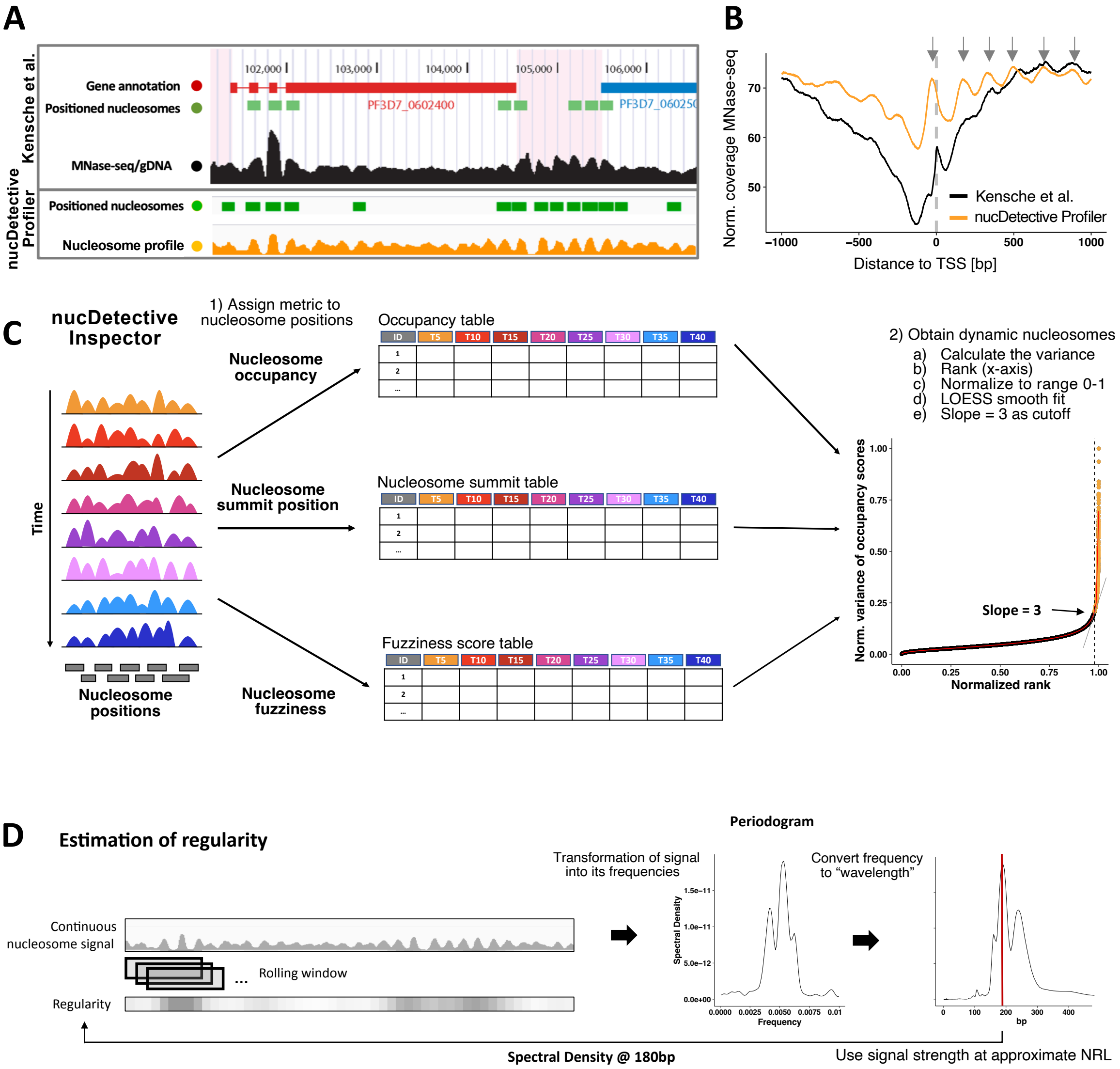


**Figure S1. nucDetective enables detection of dynamic nucleosome features at a high resolution**

(A) The optimized MNase-seq data analysis workflow Profiler of the nucDetective pipeline improves the resolution of nucleosome positions in *Pf*. A comparison of detected positioned nucleosomes and nucleosome coverage at T5 is shown between the originally published analysis by Kensche and colleagues (top panel) (Kensche et al. 2016) and our re-analysed data using the Profiler workflow of the nucDetective pipeline (bottom). This figure contains an edited figure from (Kensche et al. 2016).

(B) Re-analyzed nucleosome TSS meta profile (yellow) exhibits phased nucleosomes (grey arrows) downstream of the TSS and a positioned +1 nucleosome located at the TSS. For comparison, the results of the original analysis by Kensche and colleagues (black) are shown (Kensche et al. 2016).

(C) Scheme outlining the Inspector workflow of the nucDetective pipeline to call nucleosomes with a change in occupancy, fuzziness or position shift over time. The analysis method of the different categories follows a common procedure: First, a score is assigned for each sample (here time point) to each nucleosome position. In case of position shifts, the exact dyad position

at each timepoint is computed by loading the coverage track at the referen.ce position, fitting a smooth curve and determining the summit position. In a second step, the resulting score matrix is used to calculate the variance for each nucleosome position over all time points. The resulting variance is normalized to a range between 0 and 1 (y-axis) and plotted against the ranks normalized by the total number of nucleosome positions (x-axis). A LOESS smoothing curve is fitted (red line), and the slope of this curve is used to determine a cutoff (grey line). Here a slope cutoff of 3 was used (dashed line). Nucleosomes with a higher variance (yellow dots) are considered to indicate a change in the respective feature across all samples.
(D) Scheme outlining the regularity estimation process within the nucDetective pipeline. The nucleosome coverage profile is split into rolling windows. For each window a periodogram is computed which transforms the signal into its frequencies and assigns a portion of the observed signal to each frequency, referred to as the spectral density. The frequencies are then converted into spatial periods. The spectral density at the approximate Nucleosome Repeat Length (NRL, here 180 bp) serves as a measure of regularity for that period, which is mapped back to the original nucleosome signal window.

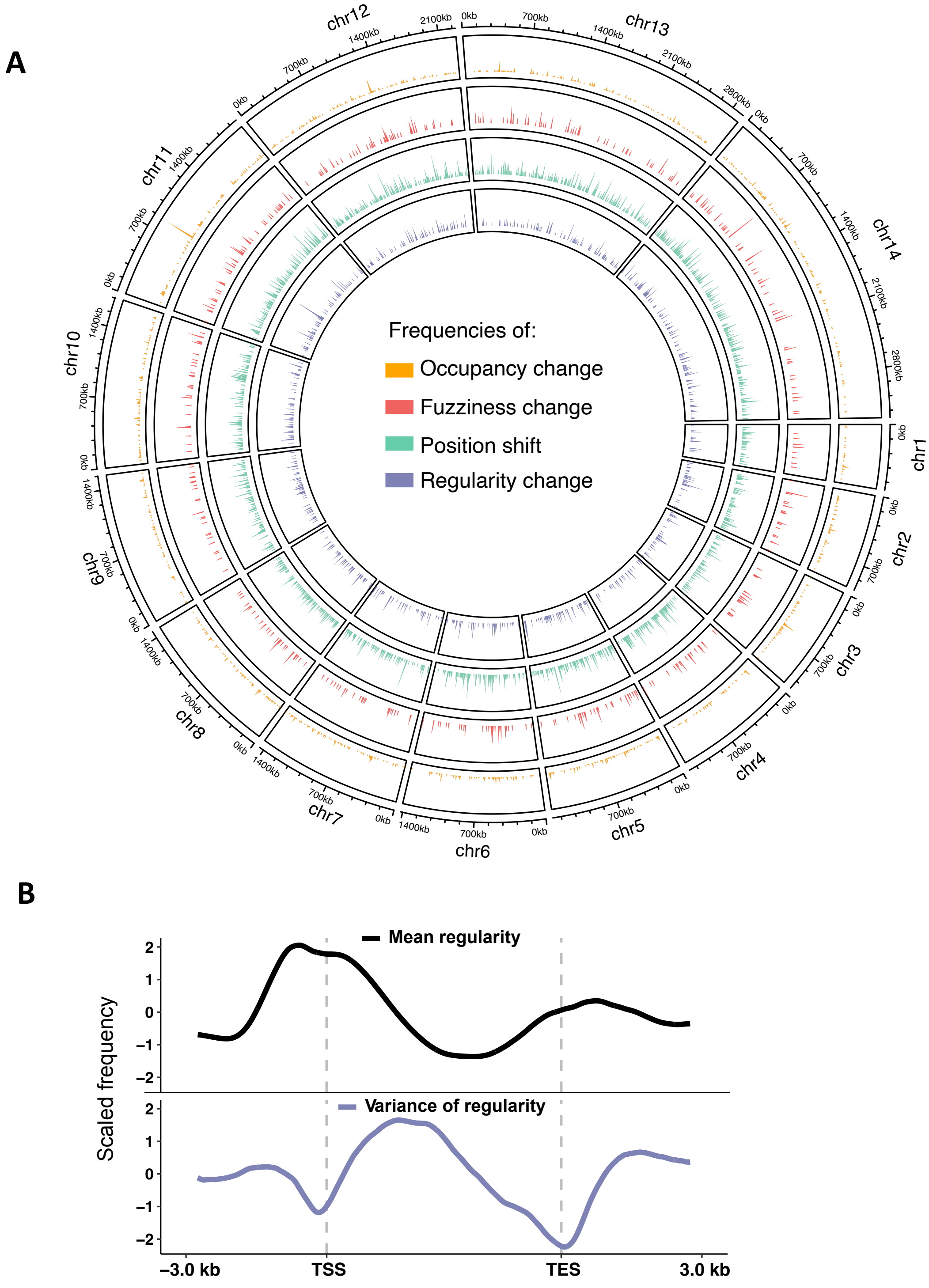


**Figure S2. Global overview of dynamic nucleosomes**

(A) Dynamic nucleosomes are evenly distributed across the entire genome on a global scale. Frequencies of nucleosomes showing occupancy (yellow), fuzziness (red), position (green) and regularity (blue) changes in 10 kb bins is depicted across the whole genome.

(B) Nucleosomes display regular spacing at the TSS and changes of regularity during the IDC are primarily observed in the gene body. The meta profile of centered and scaled mean regularity (black) and variance of regularity (blue) is plotted over length scaled gene regions. Regularity is derived from the log10 spectral power at the period of 180 bp. TES = Transcription End Site

| Cluster | Top 3 motifs | p-value | % Targets | % Back-ground | Best matches | Score | Motif name |
|---|---|---|---|---|---|---|---|
| 1 | 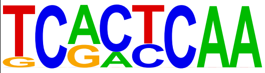 | 1e-6 | 6.13 | 0.77 | 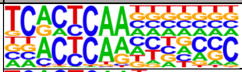 | 0.66 | PF13_0097_Secondary_motif_3 |
| | | | | | 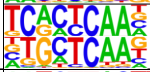 | 0.63 | PF13_0097_Secondary_motif_1 |
| | 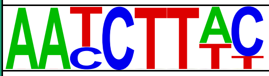 | 1e-5 | 10.43 | 2.82 | 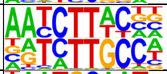 | 0.6 | PF10_0075_D2 |
| | 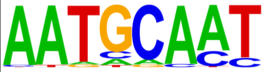 | 1e-5 | 8.59 | 1.97 | 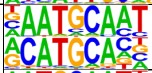 | 0.65 | PF14_0633_Secondary__motif_1 |
| | | | | | 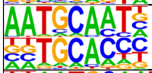 | 0.63 | PF10_0075_D3_Secondary_motif_4 |
| | | | | | 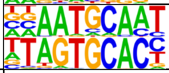 | 0.61 | PF10_0075_D3 |
| 2 | 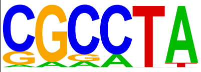 | 1e-6 | 33.96 | 13.86 | 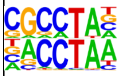 | 0.74 | PF13_0267_Secondary_motif_2 |
| | | | | | 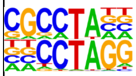 | 0.69 | PFF0670w_D2_Secondary_motif_1 |
| | 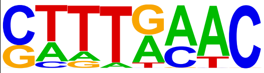 | 1e-6 | 29.25 | 11.23 | 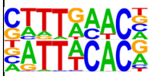 | 0.69 | PFD0985w_D2_Secondary_motif_1 |
| | | | | | 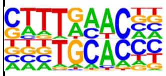 | 0.65 | PF10_0075_D3_Secondary_motif_4 |
| | | | | | 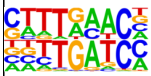 | 0.63 | PF13_0097_Secondary_motif_2 |
| | 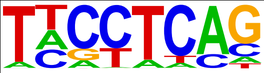 | 1e-5 | 12.26 | 2.54 | 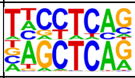 | 0.66 | PF13_0097 |
| 3 | 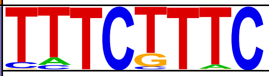 | 1e-7 | 18.21 | 8.86 | Novel motif | | |
| | 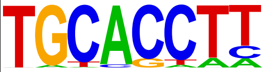 | 1e-6 | 11.27 | 4.52 | 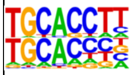 | 0.87 | PF10_0075_D3_Secondary_motif_4 |
| | | | | | 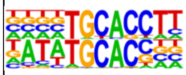 | 0.77 | PF10_0075_D3_Secondary_motif_3 |
| | | | | | 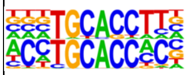 | 0.74 | PFF0200c_D1_Secondary_motif_1 |
| | 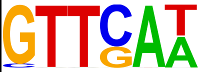 | 1e-5 | 25.43 | 15.97 | 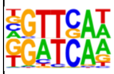 | 0.69 | PF13_0097_Secondary_motif_2 |
| | | | | | 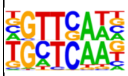 | 0.68 | PF13_0097_Secondary_motif_1 |
| | | | | | 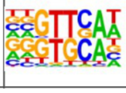 | 0.64 | PF10_0075_D3_Secondary_motif_4 |

| | | | | | | | |
|---|---|---|---|---|---|---|---|
| 4 | 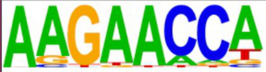 | 1e-7 | 19.71 | 9.73 | 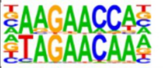 | 0.64 | PF11_0404_D1 |
| | 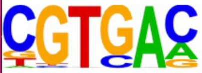 | 1e-5 | 6.47 | 1.98 | 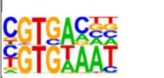 | 0.64 | PFD0985w_D2_Se-condary_motif_1 |
| | | | | | 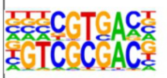 | 0.6 | PF10_0075_D1_Se-condary_motif_3 |
| | 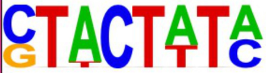 | 1e-5 | 10.00 | 4.10 | 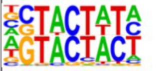 | 0.6 | PFL1085w_Secondary_mo-tif_1 |
| 5 | 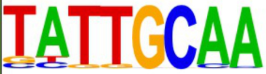 | 1e-6 | 14.16 | 3.26 | 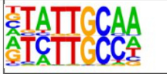 | 0.74 | PF10_0075_D2 |
| | | | | | 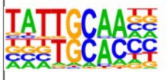 | 0.65 | PF10_0075_D3_Se-condary_motif_4 |
| | | | | | 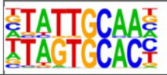 | 0.65 | PF10_0075_D3 |
| | 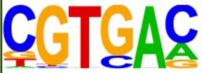 | 1e-4 | 29.20 | 13.80 | 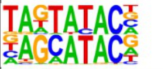 | 0.73 | PF11_0091 |
| | | | | | 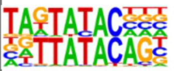 | 0.67 | PFL1075w_Secondary_mo-tif_2 |
| | | | | | 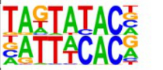 | 0.64 | PFD0985w_D2_Se-condary_motif_1 |
| | 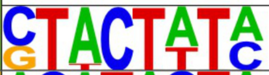 | 1e-4 | 10.62 | 2.79 | Novel motif | | |
| 6 | 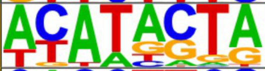 | 1e-4 | 42.74 | 25.10 | Novel motif | | |
| | 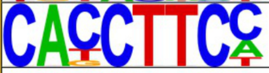 | 1e-4 | 6.45 | 0.95 | Novel motif | | |
| | 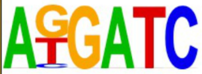 | 1e-4 | 7.26 | 1.40 | 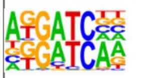 | 0.78 | PF13_0097_Se-condary_motif_2 |
| | | | | | 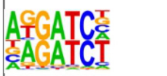 | 0.71 | PFF0670w_D2_Se-condary_motif_2 |
| | | | | | 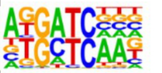 | 0.62 | PF13_0097_Se-condary_motif_1 |

**Figure S3. Distinct sequence motifs are enriched in cluster of dynamic nucleosomes**

*De novo* DNA motifs enriched in distinct clusters of nucleosome occupancy changes (cf. clustering Figure 4A). Top 3 hits of each cluster are shown along with the significance of motif enrichment (hypergeometric test) and the fraction of motifs in dynamic nucleosome cluster or random background sequences. Known *Pf* transcription factor binding motifs taken from (Campbell et al. 2010) with high similarity score (> 0.6) are shown next to it.

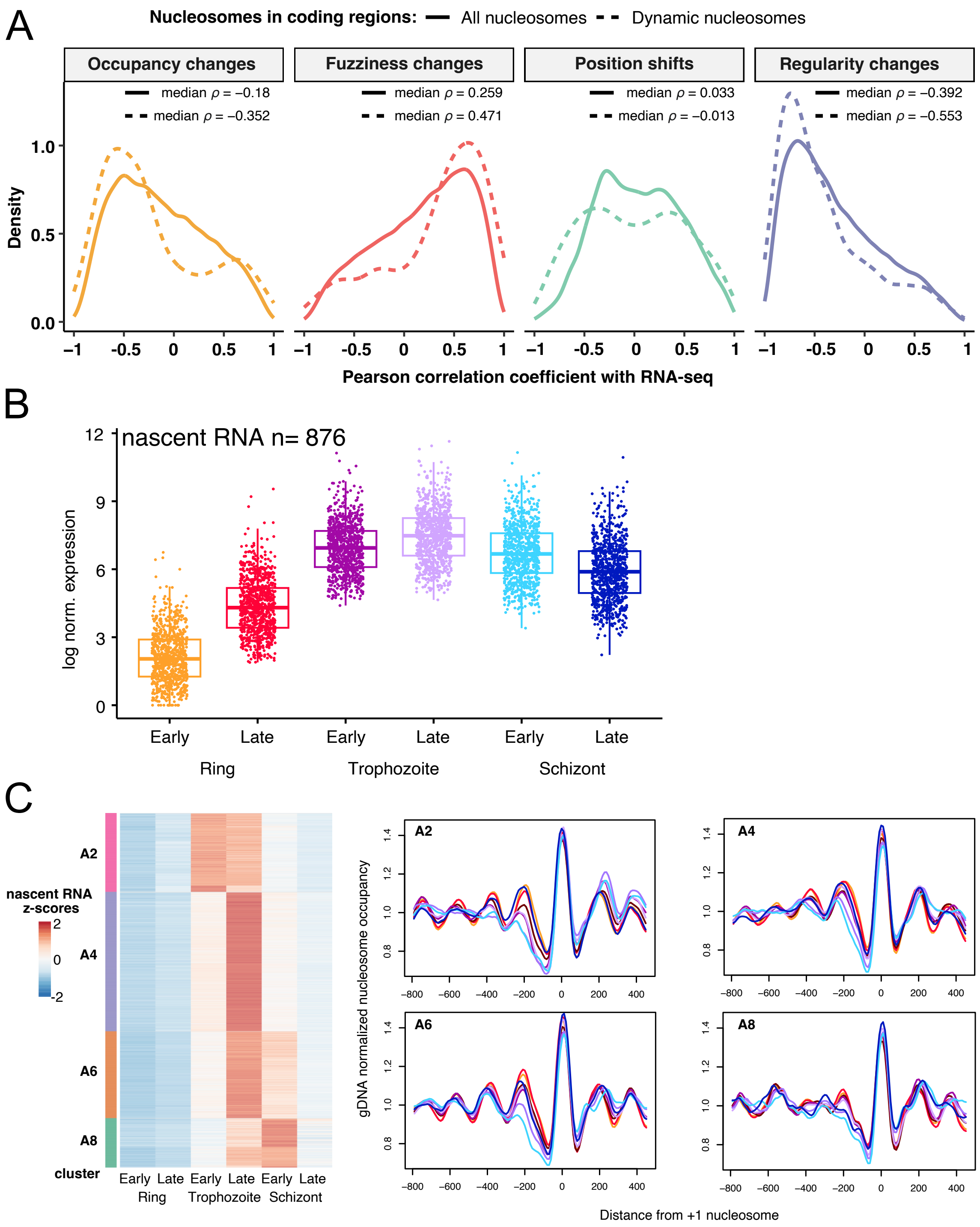


**Figure S4. Nucleosome dynamics correlate with gene transcription**

(A) Trends of nucleosome dynamics in coding regions during transcription. Linear correlations were computed for each nucleosome in coding regions, assessing the correlation between gene expression and occupancy, fuzziness, position shift and regularity over the *Pf* IDC. The density plot compares Pearson correlation coefficients for dynamic nucleosomes (dashed line) to those for all nucleosomes (solid line) in coding regions. The median Pearson correlation coefficient $\rho$ for nucleosomes with high variance (dashed line) and for all nucleosomes (solid line) are indicated.

(B) Normalized gene expression values obtained from GRO-seq data (Lu et al., 2017). The same genes as shown in Figure 5A were taken.
(C) Nucleosome features at the TSS of genes with distinct expression kinetics. Heatmap shows z-score scaled normalised nascent RNA levels measured by GRO-seq (Lu et al., 2017). Spatio-temporal expression clustering of genes as indicated on the left side was taken from Lu and colleagues. Nucleosome occupancy profiles centered at the +1 nucleosomes of clustered genes show an opening of promoter region depending on transcriptional initiation (right). Nucleosome occupancy profiles were first scaled by the underlying profile of MNase digested gDNA and then the scaled coverage profile at each time point was divided by its region median coverage value.
